# Transcriptomic analysis of melanoma cells reveals an association of α-synuclein with regulation of the inflammatory response

**DOI:** 10.1101/2023.12.23.573196

**Authors:** Santhanasabapathy Rajasekaran, Siyuan Cheng, Nithya Gajendran, Sahar Shekoohi, Liudmila Chesnokova, Xiuping Yu, Stephan N. Witt

## Abstract

The Parkinson’s disease protein, alpha-synuclein (α-syn/*SNCA*), is highly expressed in neurons and melanomas. The goal of this study was to reveal the mechanism(s) of α-syn’s involvement in melanoma pathogenesis. To decipher the genes and pathways affected by α-syn, we conducted an RNA sequencing analysis of human SK-MEL-28 cells and several SK-MEL-28 *SNCA*-KO clones. We identified 1098 significantly up-regulated genes and 660 significantly down-regulated genes. Several of the upregulated genes are related to the immune system, i.e., the inflammatory response and the matrisome. We validated five upregulated genes (IL-1ý, SAA1, IGFBP5, CXCL8, and CXCL10) by RT-qPCR and detected IGFBP5 and IL-1β in spent media of control and *SNCA*-KO cells. The levels of each of these secreted proteins were significantly higher in the spent media of the *SNCA*-KO clones than control cells. We suggest that the loss of α-syn expression unleashes chemokine/cytokine secretion, which could help melanoma cells evade the immune system.

## Introduction

Melanoma is an aggressive cancer of melanocytes, which are the pigment producing cells in the body^1^. Parkinson’s disease is a progressive neurodegenerative movement disorder characterized by the loss of dopaminergic neurons ^2,3^. Several epidemiological studies have revealed a co-occurrence of PD and melanoma, and the co-occurrence is reciprocal in that melanoma patients have a higher risk of developing PD^4,5^, and PD patients have a higher risk of developing melanoma^6–8^. The possible mechanisms of the co-occurrence, such as defects in cellular detoxification, melanin biosynthesis, oxidative stress response, and endolysosomal trafficking, and associated genes, such as Parkin, *SNCA*, LRRK2, DJ-1, *TRPM7*, *p53*, have been discussed in detail elsewhere^9,10^. Our focus here is *SNCA*.

There are curious similarities between melanocytes and dopaminergic (DA) neurons which made us focus on *SNCA*, which codes for the protein α-synuclein (α-syn). Because melanocytes and DA neurons are derived from the neural crest, they have some similar properties: both cell types express α-syn, both express enzymes that hydroxylate tyrosine (tyrosinase and tyrosine hydroxylase), both synthesize pigment (melanin and neuromelanin), pigment protects against oxidizing intermediates in each cell type, and α-syn inhibits tyrosinase which can decrease the amount of pigment in melanocytes^11^. Cytotoxic aggregated and amyloid forms of α-syn have been implicated as the causative agent in idiopathic PD^12,13^, whereas alpha-syn inhibits melanin synthesis, which could make melanocytes with high levels of α-syn susceptible to UV-radiation. Thus, in PD α-syn causes cell death, whereas in melanoma, it, along with driver mutations, leads to uncontrolled growth and metastasis.

α-Syn is a small intrinsically unfolded protein that is highly expressed in neurons, as well as other tissues such as red blood cells^14^, retinal cells^15^, and in melanoma cells^16^. α-Syn localizes to nerve terminals where synaptic vesicles are docked^17^; it accelerates the kinetics of individual exocytotic events^18^; it promotes endocytosis and vesicular trafficking in a variety of cells^19,20^; and it binds histones^21^, DNA^22^, and may control specific gene transcription by modulating epigenetic regulatory functions^17^.

We suggest that α-syn has two roles in melanoma: it can initiate pathology and then propagate it. As the initiator, by inhibiting melanin biosynthesis in melanocytes, α-syn expression increases the probability of UV-damage. UV radiation can damage the DNA, creating mutations such as BRAFV600E, which leads to uncontrolled growth and metastasis. As the propagator, that is, as a pro-survival factor, α-syn, by unknown mechanisms, promotes the proliferation of B16 murine melanoma cells^23^ and human SH-SY5Y neuroblastoma cells^24^ and promotes the motility of SH-SY5Y cells and SK-MEL-28 human melanoma cells^25^. To try to uncover the mechanisms by which α-syn might be pro-survival to melanoma, we conducted a comparative RNA-seq analysis of human SK-MEL-28 melanoma cells and several SK-MEL-28 *SNCA*-KO clones to uncover genes/pathways influenced by α-syn. Based on the findings reported herein, we suggest that α-syn may also play a role in melanoma immune evasion.

## Results

### Gene Ontology (GO) and KEGG enrichment analysis of differentially expressed genes (DEGs)

To decipher the genes/pathways affected by α-syn (*SNCA*), we conducted a comparative RNA-seq analysis of human SK-MEL-28 control cells and three SK-MEL-28 *SNCA*-KO clones with homozygous *SNCA*-knockouts (KOs) (KO6, KO8, and KO9) and one hemizygous *SNCA*-KO (KO3) (Fig. S1). The analysis yielded the identification of DEGs up- or downregulated in response to knocking out *SNCA*. Figure 1 shows a volcano plot of the up- and down-regulated genes. Overall, there were 1098 up-regulated genes and 660 down-regulated genes (adjusted *p* values < 0.000001 and log_2_FC >1 or < -1). Table 1 lists the top 15 up- and top 15 down-regulated genes. Up-regulated transcripts included *IGFBP5*, *PI3*, *MAGEL2*, *FPR3*, *IL-1ý*, *AXL*, and *CXCL10*, and down-regulated transcripts included *SLC12A7 (KCC4)* and *GALNT14*.

**Fig. 1.**
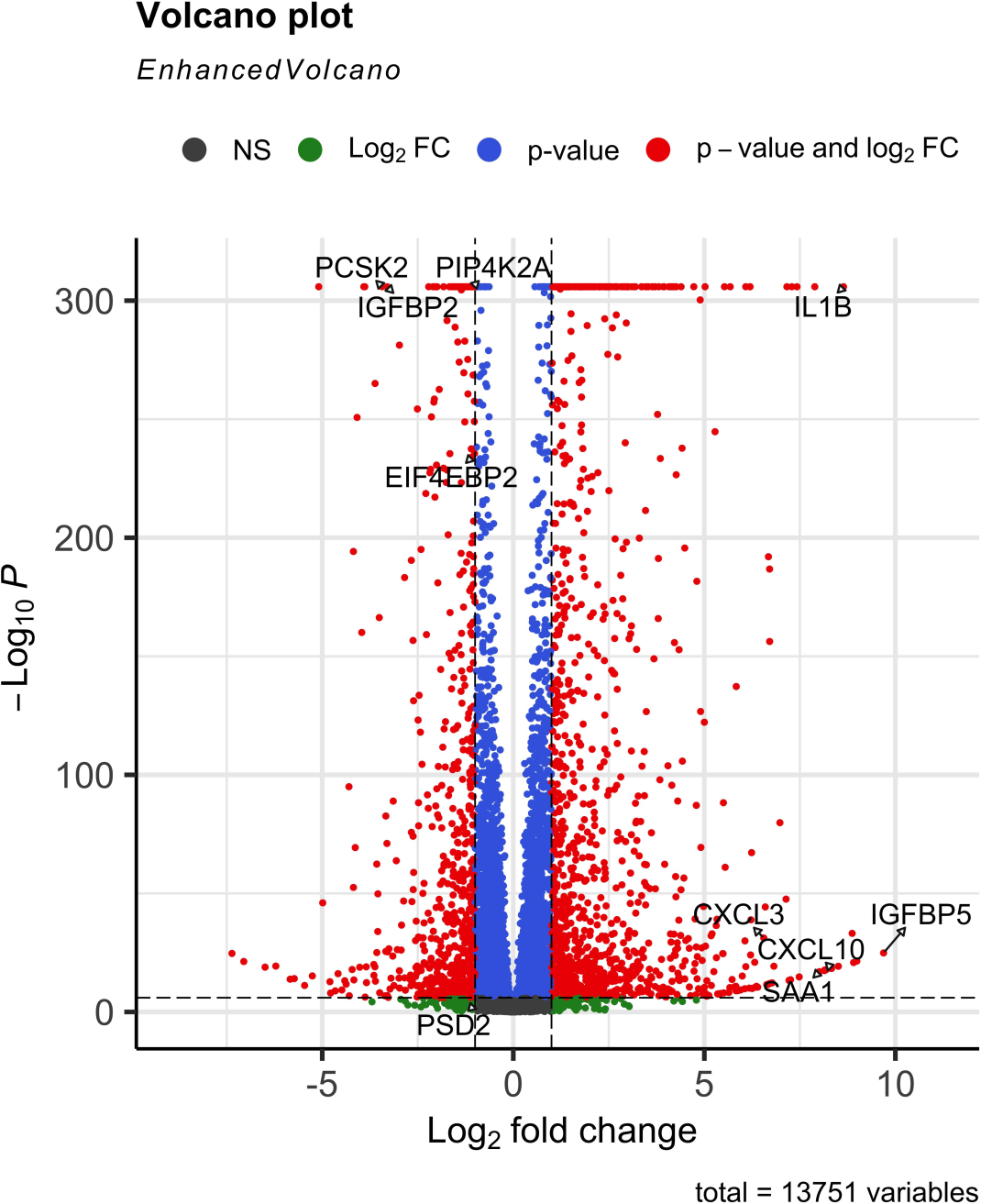
Volcano plot. The bioinformatics analysis of RNA-seq results identified significant differentially expressed (DE) genes and signaling pathways. The volcano plot was generated to visualize DE genes. The x-axis represents the log_2_FC, and the y-axis represents the significance of gene expression in knock-out versus control samples. Increasing values of “-log_10_P” indicate increasing significance.

**Table 1.** Lists some of the genes with the largest fold change.

| **Gene** | **baseMean** | **log<sub>2</sub>FC** | **p<sub>adj</sub>** |
| --- | --- | --- | --- |
| **IGFBP5** | 206.21 | 9.69 | 1.3E-25 |
| **PI3** | 126.86 | 8.99 | 3.5E-22 |
| **MAGEL2** | 53.76 | 8.91 | 1.5E-21 |
| **FPR3** | 227.96 | 8.87 | 7.3E-34 |
| **IL1B** | 3703.62 | 8.64 | 0.00 |
| **AXL** | 91.10 | 8.51 | 5.4E-20 |
| **CXCL10** | 35.98 | 8.33 | 4.7E-19 |
| **IL36RN** | 72.90 | 8.19 | 1.4E-18 |
| **SAA2** | 70.95 | 8.15 | 2.1E-18 |
| **CSF3** | 31.65 | 8.14 | 3.7E-18 |
| **SAA1** | 43.26 | 8.01 | 1.0E-17 |
| **MMP1** | 1418.55 | 7.89 | 0.00 |
| **PPBP** | 20.01 | 7.48 | 1.7E-15 |
| **SERPINB2** | 2330.32 | 7.43 | 0.00 |
| **CXCL8** | 9459.71 | 7.29 | 0.00 |
| **LCE1A** | 4.92 | -4.49 | 5.4E-10 |
| **IPO5P1** | 9.40 | -4.59 | 7.3E-17 |
| **ZDHHC22** | 4.44 | -4.66 | 1.3E-09 |
| **FOLH1** | 3.04 | -4.78 | 1.1E-08 |
| **ANKRD34C** | 7.17 | -4.80 | 1.8E-13 |
| **HUNK** | 28.52 | -4.99 | 9.4E-47 |
| **LINC00976** | 244.49 | -5.09 | 0.00 |
| **MIOX** | 9.75 | -5.26 | 6.4E-16 |
| **MAT1A** | 2.96 | -5.46 | 6.7E-12 |
| **CPE** | 9.01 | -5.73 | 8.8E-15 |
| **GALNT14** | 4.90 | -5.84 | 1.6E-14 |
| **AGXT** | 15.45 | -6.22 | 1.6E-14 |
| **C17orf77** | 12.05 | -6.50 | 1.3E-19 |
| **GREB1L** | 11.26 | -7.05 | 5.3E-22 |
| **SLC12A7** | 21.82 | -7.36 | 2.1E-25 |

The GO and KEGG enrichment analysis (http://metascape.org) identified up-regulated genes in the categories NABA matrisome associated, tube morphogenesis, extracellular morphogenesis, locomotion, interleukin-4 and interleukin-13 signaling, and the inflammatory response (Fig. 2 A, B). Down-regulated genes were in the categories herpes simplex virus 1 infection, modulation of chemical synaptic transmission, and axon guidance, among others (Fig. S2 A, B). Our focus in this report was the immunological aspects of *SNCA*. To that end, heatmaps of genes involved in the inflammatory response and the NABA matrisome-associated are shown in Figs. 3A and B, respectively. Several of these genes were validated, as follows.

**Fig. 2.**
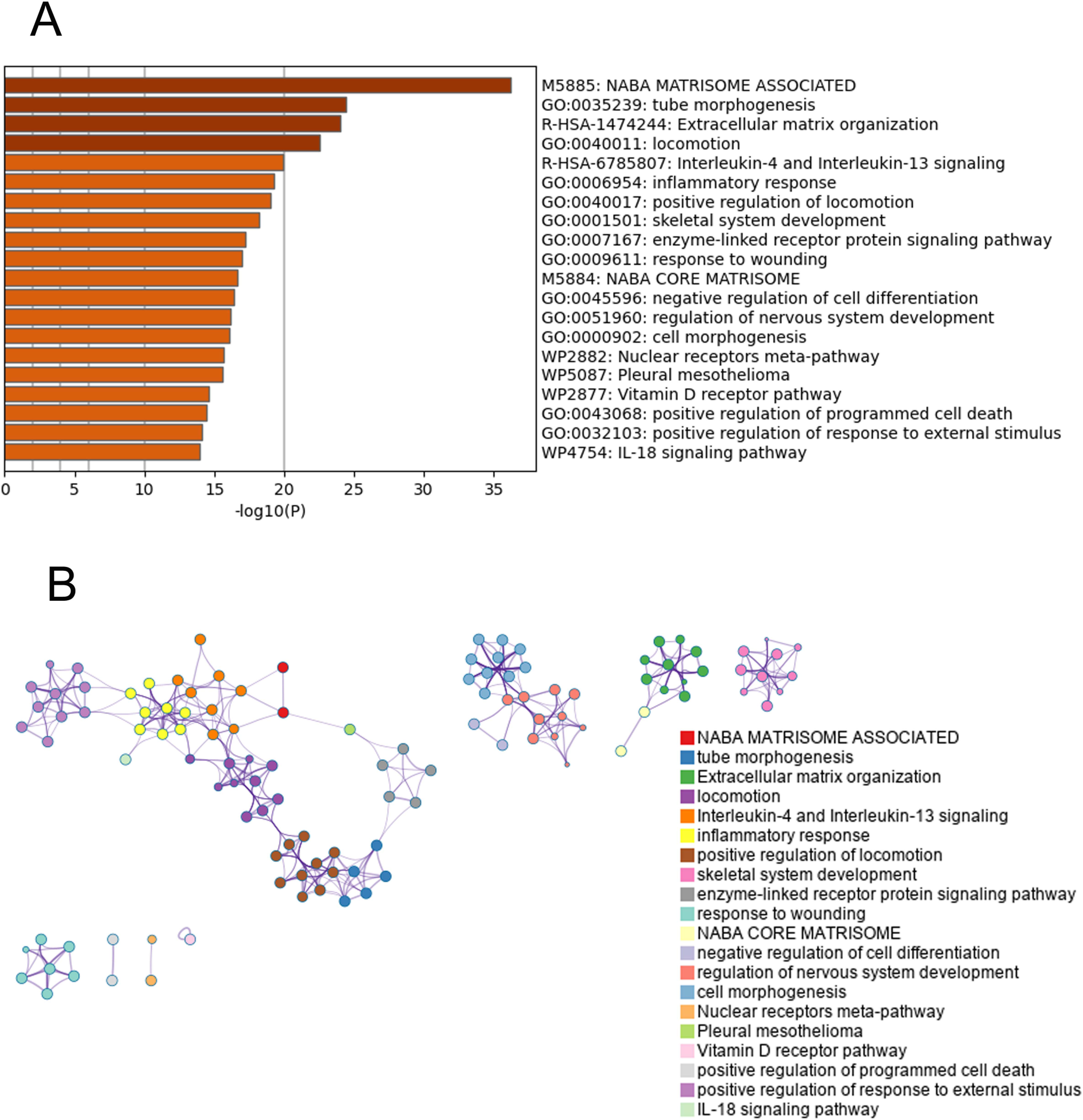
GO enrichment analysis. (**A**) Gene ontology enrichment analysis shows 20 enrichment clusters of up-regulated in *SNCA*-KO clones. (**B**) Enrichment network visualization of up-regulated genes in *SNCA*-KO clones. Both analyses conducted at http://metascape.org.

**Fig. 3.**
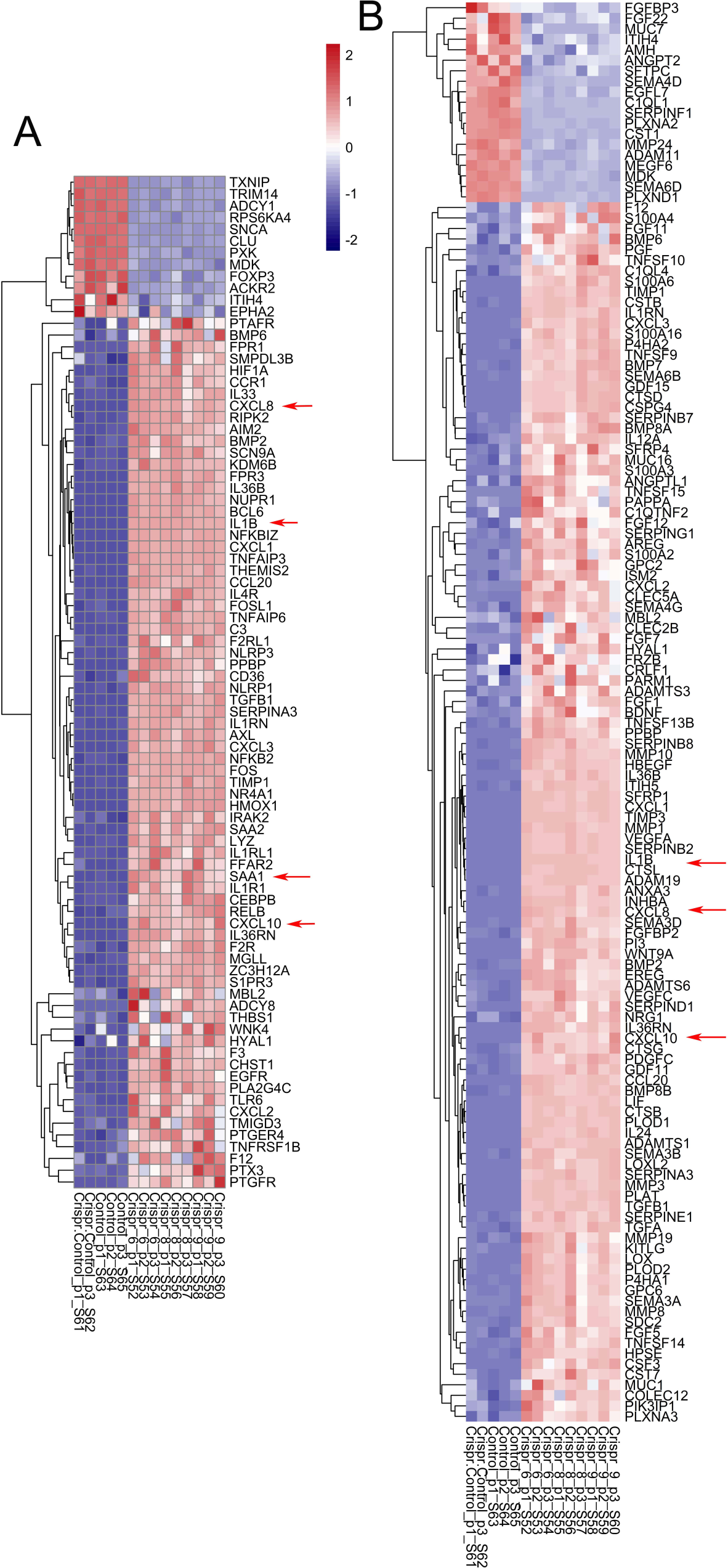
Heatmap. Bioinformatics analysis of the RNA-seq results identified significant differentially expressed (DE) genes and signaling pathways. Heatmap (**A**) was generated to visualize the significant DE genes that belong to GO category “regulation of inflammatory response.” Heatmap (**B**) show significant DE genes that belong to GO category “NABA Matrisome Associated.”

### Validation of RNA–seq data by qPCR

We validated five of the up-regulated genes, four of which are indicated in the heatmaps (Fig. 3), and two of the down-regulated genes by PCR methods (Table 2). In some cases, we validated changes in transcript levels in the human melanoma cell line WM983B, which expresses α-syn, in two WM983-B *SNCA*-KO clones, in the human neuroblastoma cell line SH-SY5Y, which expresses very low levels of α-syn, and the SH-SY5Y α-syn overexpressing line (SH/αS)^26^ (Table 3). Additionally, we detected secreted IL-1ý and IGFBP5 in the spent media of SK-MEL-28 and WM983B cells. Using these other cell lines broadened the scope of our RNA-seq results, as shown below.

**Table 2.**
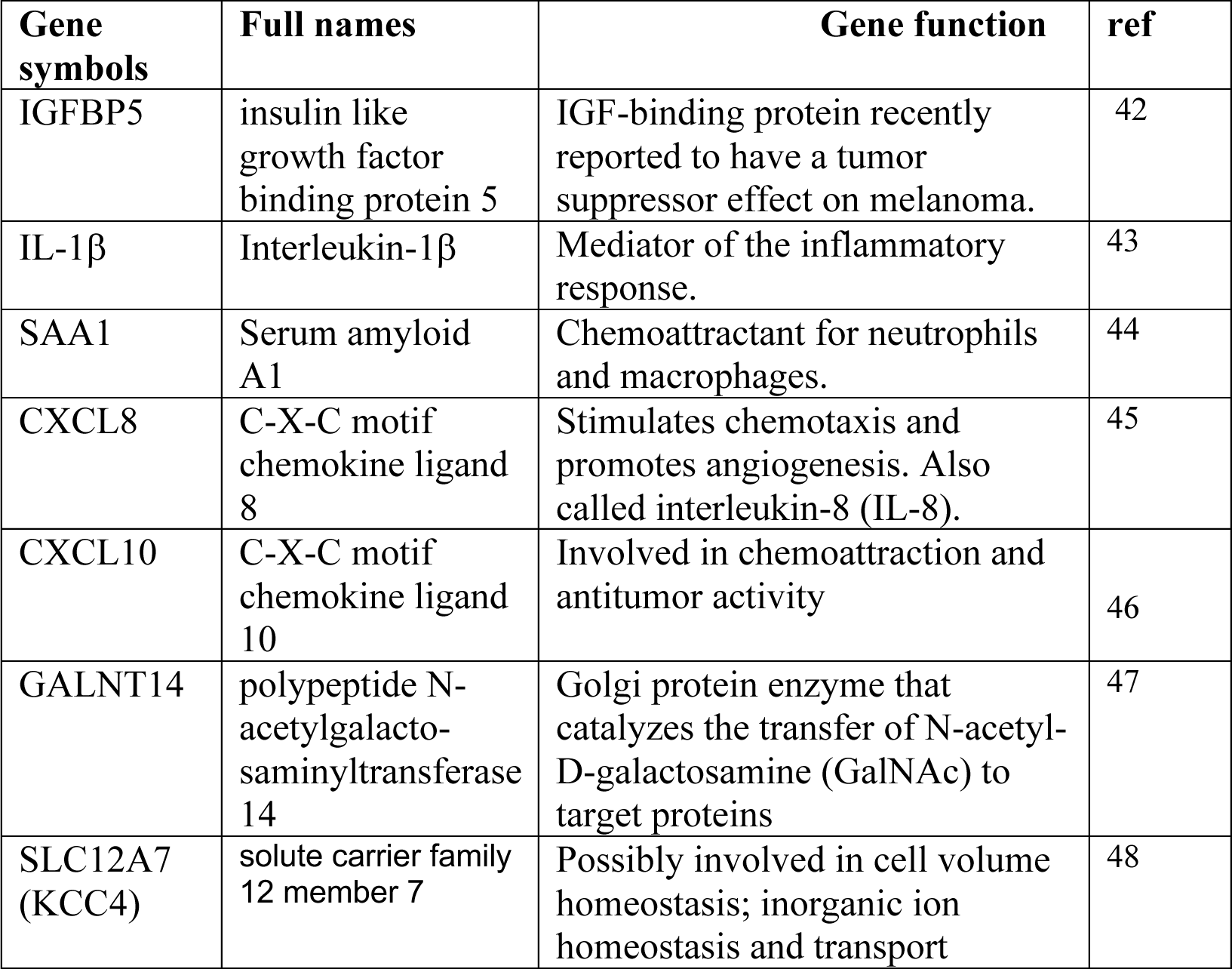
Genes selected for validation.

**Table 3.** Cell lines used in this study. α-Syn expression was tested by Western blotting and qPCR.

| **Cell line** | **Comments** | **α-syn (Y, N)** | **ref** |
| --- | --- | --- | --- |
| SK-MEL-28 | This cell line was isolated from skin tissue from a 51-year-old, male with malignant melanoma. | Y |  |
| SK-MEL-28 *SNCA*-KO clones KO8, KO9 | *SNCA* knocked out in SK-MEL-28 cells using CRISPR/Cas9 | N | 27 |
| SK-MEL-28 *SNCA*-KI clones KI8, KI9 | *SNCA* cDNA was re-introduced into the KO clones using lenti virus | Y | 27 |
| WM983B | This cell line was established from an inguinal node metastatic site in a 54-year-old male patient. | Y |  |
| WM983B *SNCA*-KO clones 1 and 2 | *SNCA* knocked out in WM983B cells using CRISPR/Cas9 | N | This study |
| SH-SY5Y | Cell line derived from a neuroblastoma from a 4-year-old child | N | 49 |
| SH/aS | SH-SY5Y stably expressing a-syn | Y | 49 |

qPCR was conducted to assess transcript levels in control and KOs for the up-regulated genes IL-1ý, IGFBP5, SAA1, CXCL8, and CXCL10 (Figs. 1, 3; Table 2). Transcript levels were also evaluated for two down-regulated genes (GALNT14 and SLC12A7) (Table 2, Fig. S3). Western blotting confirmed that SK-MEL-28 control cells express α-syn whereas the *SNCA*-KO clones KO8 and KO9 do not (Fig. 4A). qPCR analysis revealed that the IL-1β, IGFBP5, SAA1, CXCL8, and CXCL10 transcripts were all higher in the each of the two KO clones compared to control cells, with eight of ten comparisons showing significant increases between KO and control Fig. 4B). Compared to the controls, the mean transcript levels for KO8 and KO9 increased by 108%, 2,037%, 494%, 263% and 314% for IL-1β, IGFBP5, SAA1, CXCL8, and CXCL1, respectively.

**Fig. 4.**
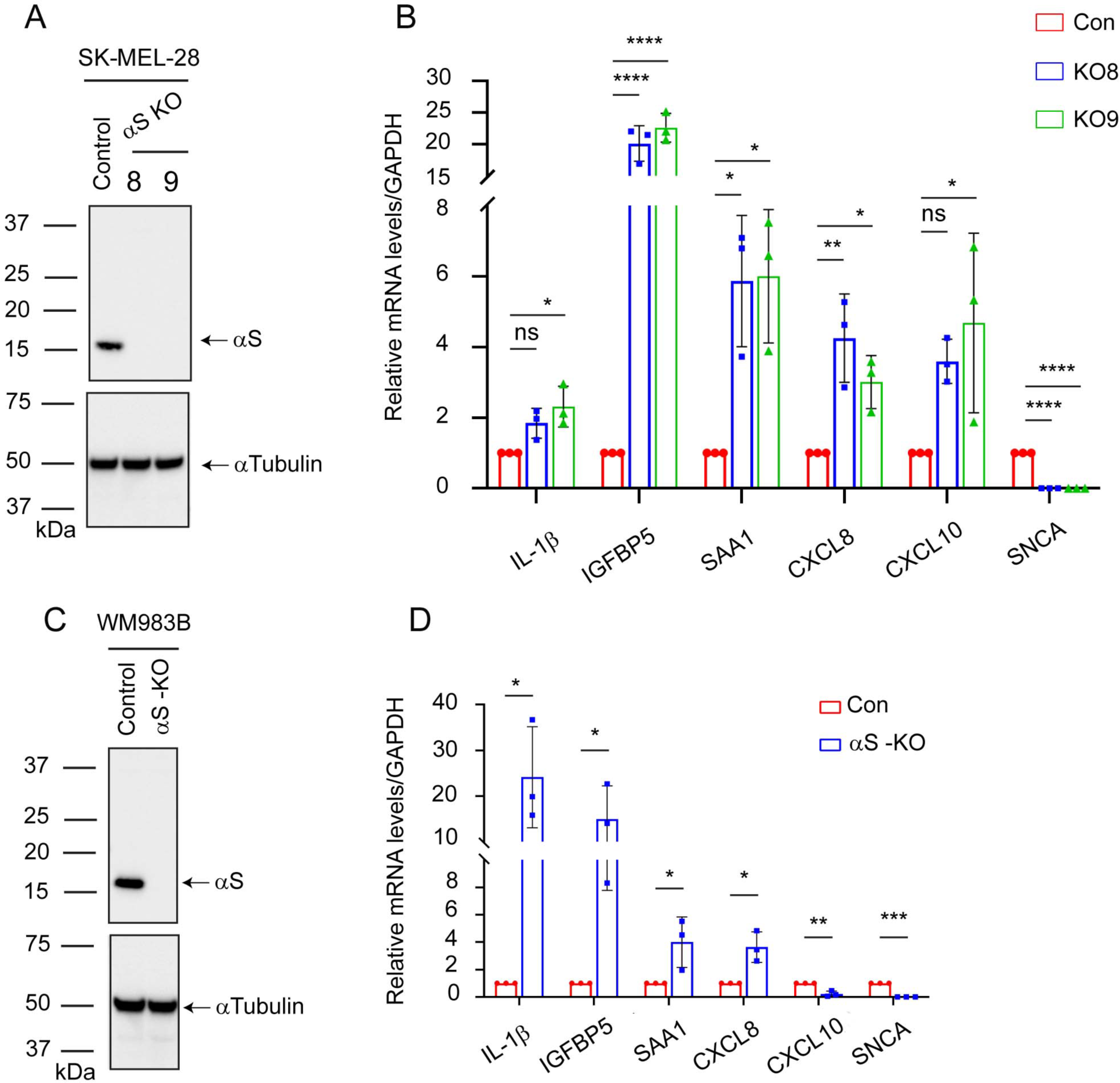
RT-qPCR validation of RNA-seq data using SK-MEL-28 and WM983B cells. (**A**) Representative western blots of α-syn and α-tubulin in lysates of the control, KO8, and KO9 cells cultured *in vitro.* (**B**) RT-qPCR analyses of differentially expressed genes in the control and KO cells. Relative mRNA levels in fold-change of *IL-1β, IGFBP5*, *SAA1, CXCL8, CXCL10,* and *SNCA* normalized to housekeeping gene *GAPDH*. Relative expression was calculated using the comparative CT method (2^−ΔΔCT^). Triplicates were used per biological sample. Data are mean ± s.d. A one-way ANOVA with Dunnett post hoc test was used to determine p values (n = 3). *, *p* :: 0.05; **, *p* :: 0.01; ****, *p* :: 0.0001. (**C**) Representative western blot of α-syn and α-tubulin in lysates of the control and α-syn KO WM983B cells cultured *in vitro*. (**D**) RT-qPCR results show gene expression in control and α-syn KO cells on selected transcripts. The comparative CT method (2^−ΔΔCT^) was adopted to calculate the relative mRNA levels of respective genes normalized to the housekeeping gene *GAPDH* and represented in graphs. Transcripts analyzed are *IL-1β*, *IGFBP5, SAA1, CXCL8*, *CXCL10*, and *SNCA*. Data are mean ± s.d. Statistical analysis was performed by using a one-tailed Student’s t-test with unequal variances (n=3). *, *p* < 0.05, ***, *p* < 0.001. Uncropped blots are shown in Fig. S4.

We also tested whether the transcripts for IGFBP5, IL-1ý, SAA1, CXCL8, and CXCL10 were up-regulated in a *SNCA*-KO clone of the WM983B human cutaneous melanoma cell line. Figure 4C shows western blot analysis of lysates of WM983B control cells and the *SNCA*-KO clone; the KO clone is devoid of α-syn expression. qPCR analysis revealed significant increases in the IL-1ý, IGFBP5, SAA1, and CXCL8 transcripts and a significant decrease in the CXCL10 transcript (Fig. 4D). Compared to the control, the mean transcript levels in the *SNCA*-KO clone increased by 2,313%, 1402%, 300% and 263% for IL-1β, IGFBP5, SAA1, CXCL8, and decreased 78% for CXCL10.

### Knocking out α-syn expression increases the secretion of IL-1β and IGFBP5

Given the higher levels of IGFBP5 and IL-1β transcripts in the *SNCA*-KO clones relative to the control cells, we asked whether the KO clones secrete more IGFBP5 and IL-1β than control cells. To detect these two proteins, we used the human IL-1 β/IL-1F2 Quantikine ELISA Kit and human IGFBP5-5 Duo set Quantikine ELISA kit. We also used SK-MEL-28 *SNCA*-KI clones, where α-syn was reintroduced into the respective KO clone via lentivirus (Fig. 5A). We detected on average 20 pg/ml of IL-1β from SK-MEL-28 control cells, and this value increased to 45 pg/ml and 60 pg/ml (*p* = 0.0023) for KO8 and KO9 (Fig. 5B), respectively. Clones KI8 and KI9 secreted approximately 10 pg/ml. Using the same ELISA assay, we tested for IGFBP5 secretion in the various SK-MEL-28 cells. Control and KI clones secreted 5 ng/ml of IGFBP5, whereas KO clones secreted approximately twice as much IGFBP5, i.e., KO8 and KO9 secreted 10 ng/ml (*p* = 0.0118) and 11 ng/ml (*p* = 0.0046) (Fig. 5C), respectively.

**Fig. 5.**
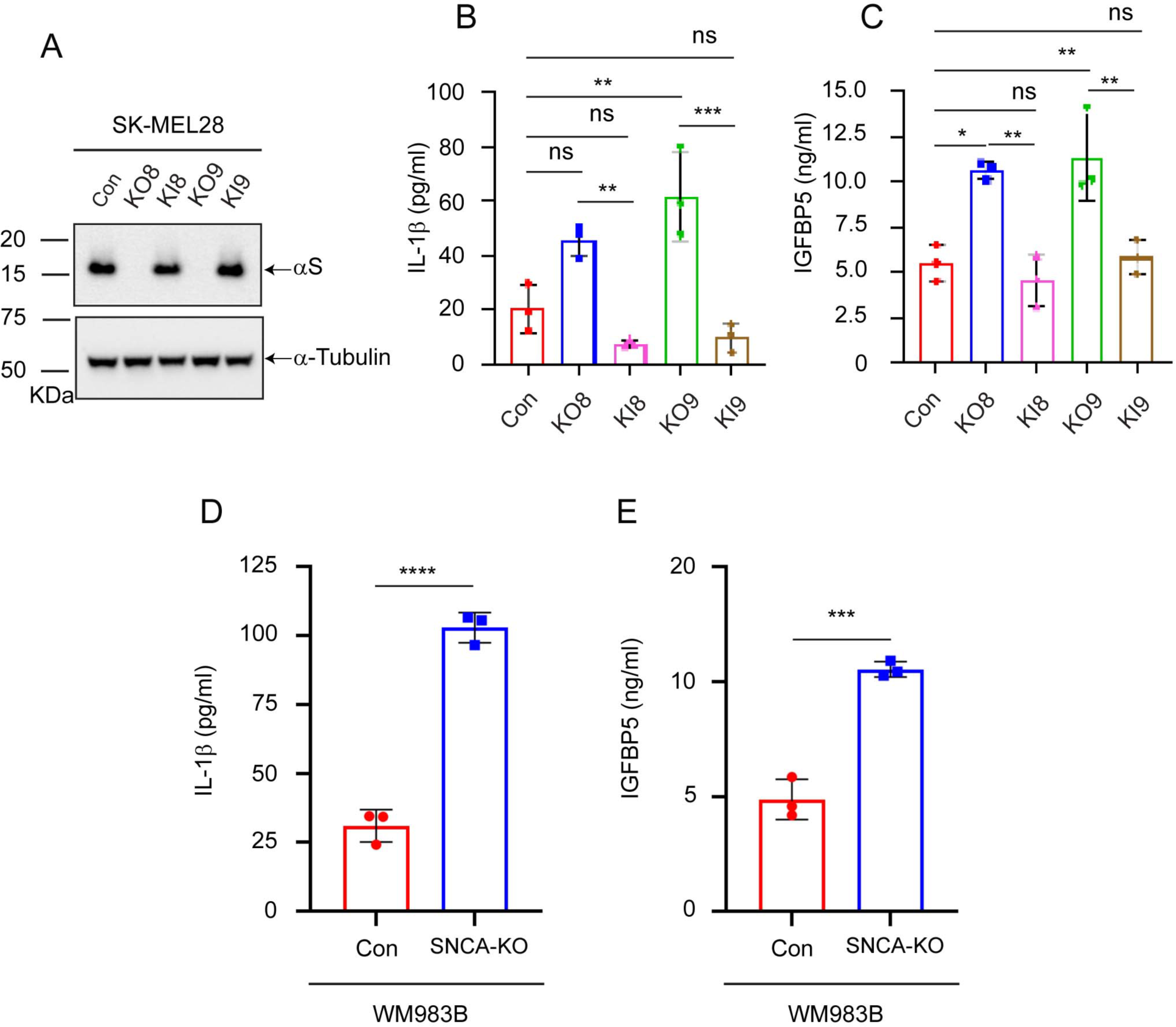
Loss of α-syn expression increases IL-1β and IGFBP-5 secretion in SK-MEL-28 and WM983B cells. (**A**) Representative western blot of α-syn and α-tubulin in lysates of SK-MEL-28 control, *SNCA*-KO, and *SNCA*-KI cells. ELISA analysis of secreted IL-1β (**B**) and IGFBP5 (**C**) levels from the culture medium of control, *SNCA*-KO (KO8 and KO9) and *SNCA*-KI (KI8 and KI9) cells cultured *in vitro.* Two technical replicates were used per biological sample. Data are mean ± s.d. P-values were determined using a one-way ANOVA with a Bonferonni correction for multiple comparisons (n=3). ELISA analysis of secreted IL-1β (**D**) and IGFBP5 (**E**) levels from the culture medium of WM983B control, *SNCA*-KO cells cultured *in vitro.* Two technical replicates were used per biological sample. Data are mean ± s.d. P-values were determined using a one-tailed Student’s t-test with equal variances (n=3). ***, *p* < 0.001, ****, *p* < 0.0001. Uncropped blots are shown in Fig. S4.

We also tested for secretion of IL-1ý and IGFBP5 from the WM983B control and KO cells. Using the ELISA assay, we found that the WM983B *SNCA*-KO clone secreted approximately 300% more IL-1ý than the control cells (*p* = 5.7 x 10^-5^). Similarly, the WM983B *SNCA*-KO clone secreted approximately 120% more IGFBP5 than the control cells (*p* = 2.3 x 10^-4^) (Fig. 5D). Collectively, except for CXCL10, the above results agree with the RNA-seq transcriptomic analysis. Additionally, we demonstrated robust secretion of IL-1β and IGFBP5 in the WM983B *SNCA*-KO clone compared to control cells.

## Discussion

This work was undertaken to determine which genes are differentially expressed in human cutaneous melanoma cells in which the *SNCA* gene was knocked out. Given that we have previously reported that SK-MEL-28 *SNCA*-KO cells are defective in migration, invasion, motility, and growth^25,27^, we anticipated these pathways, and many others, would be enriched with differentially expressed genes (Figs. 2A, B and S2A, B). Overall, the results provided herein are a blueprint for future studies.

One enrichment cluster of the down-regulated genes, i.e., “L1CAM interactions,” is of particular interest (Fig. S2). We recently reported that SK-MEL-28 *SNCA*-KO clones have significantly lower levels of the neuronal adhesion protein L1CAM compared to control cells^25^, and, reciprocally, increasing the level of α-syn expression in the neuroblastoma cell line SH-SY5Y significantly increased both L1CAM and single-cell motility. Because L1CAM, which is a signal transducer, is involved in cell signaling^28^, a decrease in the level of L1CAM in KO clones could certainly disrupt or eliminate L1CAM interactions.

Another pathway in our GO analysis of the RNA-seq results that is consistent with our previous work is the upregulation of genes in the category “positive regulation of programmed cell death” (Fig. 2A, B). We previously reported that SK-MEL-28 *SNCA*-KO xenografts exhibit increased levels of both ferric iron and ferritin relative to controls, and we tested whether the xenografts exhibited increased apoptosis^27^. Xenograft sections were analyzed for DNA fragmentation using the TdT-mediated, dUTP-biotin nick-end labeling (TUNEL) assay. We found that, on average, the sections from the KO xenografts had two-to three-fold higher amounts of DNA fragmentation than SK-MEL-28 control cells (which expressed α-syn). Collectively, *SNCA*-KO xenografts had higher levels of apoptotic markers than the control xenografts, which had implied an upregulation of programmed cell death genes, which we have verified herein.

Two GO categories that were upregulated in the *SNCA*-KO cells relative to control cells were the “inflammatory response” and “locomotion (Fig. 2A, B).” Some of the genes in these categories were validated by RT-PCR and ELISA. When we refer to locomotion it does not mean migration and metastasis of melanoma cells; instead, we suggest that locomotion refers to the movement or infiltration of immune cells (T cells, neutrophils, and macrophages) in response to the secretion of the cytokines /chemokines (CXCL8, CXCL-10, IL-1ý, and possibly others that we did not validate) by the *SNCA*-KO tumor cells. The possible roles of these secreted factors are as follows.

CXCL8 secreted by melanoma cells promotes tumor growth and angiogenesis^29^, and elevated CXCL8 levels in the serum of melanoma patients positively correlates with disease progression^30^. CXCL10 binds to and activates the CXCR3 receptor, which is predominantly expressed on activated T lymphocytes (Th1) natural killer (NK) cells, inflammatory dendritic cells, macrophages, and B cells^31^. Co-expression of CXCL10 and CXCR3 predicts an increased metastatic potential and poor overall survival^32^. However, a recent study argues that increased production of CXCL10 in the tumor microenvironment by immune cells is a positive prognostic factor for response to immunotherapy^33^ suggesting the source of CXCL10, target cells and the stage of cancer dictate the outcome of immune responses. IL-1β secreted by melanoma cells promotes macrophage infiltration and angiogenesis^34^. The upregulation of the IL-4 and IL-13 signaling pathways (Fig. 2A, B) suggest that these cytokines, if secreted by *SNCA*-KO cells, may lead to an increase in the Th1/Th2 imbalance in the tumor microenvironment leading to Th2 skewing which may promote tumor progression^35^. Lastly, IGFBP5 is a tumor suppressor^36^. Collectively, these findings raise the possibility that the secretion of these cytokines could decrease the anti-tumor response against *SNCA*-KO melanoma cells. Alternatively, we cannot rule out the possibility that *SNCA*-KO melanoma cells are ‘immunologically hot’ in the sense that they more efficiently recruit immune cells into the tumor microenvironment which enhances anti-tumor responses.

The matrisome is an ensemble of over 1000 genes that encode extracellular matrix (ECM) and ECM-associated proteins^37^. The core matrisome is made up of collagens, fibronectins, laminins, fibrillins, proteoglycans, and many other proteins, whereas matrisome-associated proteins fall into the categories ECM-affiliated (mucins, C-type lectins, annexins, galectins, etc.) and ECM regulators (lysyl oxidase, cathepsins, tissue inhibitors of metalloproteinases (TIMPS), transforming growth factor beta (TGFβ), bone morphogenic proteins (BMPs), Wnts, and cytokines). We discussed above how the loss of α-syn expression results in the upregulation of transcripts for cytokines and chemokines. The loss of α-syn expression also results in the upregulation of transcripts of lysyl oxidase (LOX and LOXL2), matrix metalloproteinases (MMP1,8,10,13) and their inhibitors (TIMP1, TIMP3), TGFβ, and other genes (Fig. 3B). Depending on the balance of pro- and anti-tumorigenic factors secreted by *SNCA*-KO melanoma cells, the net result will likely be immune infiltration and suppression of tumor growth or remodeling of the ECM such that tumor cells escape the ECM and proliferate. Clearly, more experimentation is required to determine how loss of α-syn expression in melanoma cells affects the immune system.

In sum, this is the first report of RNA sequencing of a human cutaneous melanoma cell line in which the gene coding for the Parkinson’s disease-associated protein, α-syn, is knocked out. We have already shown that in vitro the SK-MEL-28 *SNCA*-KO clones are less invasive and less migratory than the control cells that express α-syn, and that these KO clones grafted into SCID mice grow significantly slower than the control cells^27^. All these results suggest that knocking down the expression of α-syn might have a therapeutic benefit for melanoma patients. The issue raised by this study is whether the loss of α-syn expression in melanoma cells enhances or suppresses the immune response to such tumors.

## Materials and methods

### Cell lines and cell culture

The SK-MEL-28 was purchased from American Type Culture Collection (ATCC, Manassas, VA). SK-MEL-28 parental cells and SK-MEL-28 *SNCA*-KO clones were propagated in DMEM supplemented with 10% fetal bovine serum (FBS) and 1% penicillin–streptomycin. WM983B was purchased from Rockland Immunochemicals, Inc., PA and cultured in RPMI 1640 medium supplemented with 10% fetal bovine serum (FBS) and 1% penicillin–streptomycin. CRISPR/Cas9 genome editing was used to target *SNCA* in WM983B cells as described previously for SK-MEL-28 cells^27^ using α-syn CRISPR/Cas9 knockout plasmid (Santa Cruz Biotechnology # sc-417273-NIC). Lentivirus particles expressing human α-syn under cytomegalovirus (CMV) promoter (Applied Biological Materials, Inc, Canada) was used to re-express α-syn in SK-MEL-28 *SNCA*-KO cells as described previously^27^

### RNA sequencing

After harvesting SK-MEL-28 (controls and KOs) cells, the total RNA was extracted using the E.Z.N.A column-based total RNA kit (Omega BioTek, catalog number R6834-01) following the manufacturer’s protocol. Total RNA concentration was determined using Nanodrop spectrophotometer. Samples were then assessed for RNA integrity number (RIN) in the LSUHSC Research Core Facility. The maximum desirable score for this test is 10, and the samples we isolated had RINs > 8. The samples were shipped on dry ice to Indiana University (IU) School of Medicine Center Medical Genomics for RNA sequencing. The samples sent for sequencing were *SNCA*-KO3 (n=3), *SNCA*-KO6 (n=3), *SNCA*-KO8 (n=3), *SNCA*-KO9 (n=3), SK-Mel-28 Control (n=3), and SK-Mel-28 Crispr-Control (n=2). The “SK-Mel-28 Control” cells were purchased from ATCC. The SK-Mel-28 “Crispr-Control” cells were SK-MEL-28 cells that were sorted into single cells. A single SK-MEL-28 cell was then expanded, and this clone designated as the “Crispr-Control.” An Illumina HiSeq 4000 instrument was used for the analysis.

### Western blotting

Preparation of cell lysates, SDS/PAGE, and western blot analysis was carried out as described previously^27^. Protein samples were extracted from total cell lysates using RIPA buffer (50 mM Tris HCL, pH 7.4, 1% NP-40, 0.5% sodium deoxycholate, 0.1% SDS, 5 mM EDTA) and subjected to electrophoresis using NuPAGE™ 4 to 12%, Bis-Tris precast polyacrylamide gel (Invitrogen # NP0323BOX) and subsequently transferred to polyvinylidene difluoride (PVDF) membrane using (Trans-Blot® Turbo™ Mini PVDF Transfer Pack, Bio-Rad # 1704156). The membranes were often cut into strips and the individual strips were hybridized with indicated primary antibodies overnight at 4°C followed by incubation with respective horse radish peroxidase conjugates. Protein bands were detected using an enhanced chemiluminescence substrate (Clarity™ Western ECL Substrate, Bio-Rad #170-5060) and the images were acquired using Biorad Chemidoc-MP imaging system. Image J software was used to quantify protein levels. α-Tubulin blots were used as loading controls.

### RNA extraction, cDNA preparation, and qPCR

Total RNA was extracted from cells using E.Z.N.A column-based total RNA kit (Omega BioTek). Total RNA concentration was measured using a NanoDrop spectrophotometer (Thermo Scientific) and the purity of the samples was estimated by the OD ratios (A260/A280, ranging within 1.8–2.2). cDNA was synthesized using oligo(dT)/random primer mediated reverse transcription using the iScript cDNA Synthesis Kit (Bio-Rad) with 1 μg of DNA-free total RNA as input. Amplification was performed with Applied Biosystems TaqMan™ Gene Expression Assays with primer/probe sets for *SNCA* (Hs00240906), *SAA1* (Hs00761940), *IGFBP5* (Hs00181213), *CXCL10* (Hs00171042), *CXCL8* (Hs00174103), *IL-1β* (Hs00174097), *GAPDH* (Hs02786624). using Bio-Rad CFX384 Touch Real-Time PCR System. Data analyses were performed using Bio-Rad Software to measure the threshold cycle (CT) for each reaction. The comparative CT method (2^−ΔΔCT^) was used for the relative quantification of the target gene. Data were normalized to *GAPDH* messenger RNA (mRNA) levels and expressed as a relative fold change compared to control samples. For the qPCR analysis of *GALNT14* and *SLC12A7*, we used primer/probe sets *GALNT14* (Hs.PT.58.38663215) and *SLC12A7* (Hs.PT.58.192727890) purchased from IDT.

### ELISA

The secretion of cytokines IL-1β and IGFBP5 was quantified using human IL-1ý/IL-1F2 Quantikine ELISA Kit (DLB50, RD Systems) and Human IGFBP5 Duo set Quantikine ELISA kit (DY875, RD Systems). Briefly, when the cells reached confluence, they were washed with PBS and transferred to serum-free media. After 48 hours the media was collected and centrifuged at 3000 rpm for 5 minutes to remove the cell debris and concentrated using Amicon ultra centrifugal filter 10 kDa cutoff filters (UFC9010, Millipore Sigma). The concentrated media was subjected to the ELISA assay according to the manufacturer’s protocol. The absorbance was detected at the appropriate wavelength using (ThermoFisher Scientific) plate readers. IL-1β and IGFBP5 concentrations were calculated from the standard curve for the respective protein by plotting the mean absorbance (y-axis) against the protein concentration (x-axis).

### GO and KEGG enrichment analysis of differentially expressed genes (DEGs), RNA-seq analysis

The RNA-seq data was analyzed using R with custom codes ( https://github.com/schoo7/SCNA_KO). The “DEseq2” R package was used for the DE analysis with RNA-seq gene count matrix as input^38^. The log_2_FC >1 or < -1 and adjusted P-value < 0.05 were used as cutting off condition for selecting significant DE genes. The “clusterProfiler” package (https://bioconductor.org/packages//2.10/bioc/html/clusterProfiler.html) and “WebGestalt” (https://www.webgestalt.org/) were used for gene function enrichment analysis^39,40^. The “pheatmap” package (https://www.rdocumentation.org/packages/pheatmap/versions/1.0.12/topics/pheatmap) was used to visualize the transcript per million (TPM) values of the DE genes^41^.

### Statistical analyses

Hypothesis testing methods included a one-way analysis of variance (ANOVA) with a Dunnett post hoc test or Bonferroni correction when comparing multiple groups to control or a two-tailed Student’s t-test when comparing two groups. All data were analyzed using GraphPad Prism (version 6) software or Kaleida Graph (version 4.5) and expressed as mean ± standard deviation (s.d.) of at least three independent experiments. A *p*-value of < 0.05 was considered significant.

## Data availability

The fastg.gz files are archived at GEO.

## Code availability

The RNA-seq data was analyzed using R with custom codes ( https://github.com/schoo7/SCNA_KO).

## Acknowledgments

This study was supported by funds from the Feist-Weiller Cancer Center and the former Chancellor (Ghali) of LSU Health Sciences Center in Shreveport to S.N.W.

## Author Contributions

L.C.: validated down-regulation of GALNT14 and KCC4.

N.G. contributed to the study’s conception, experimentation, data analysis, interpretation, and validated RNA-seq by ELISA in WM983B and SK-MEL-28 cells.

S.C.: Conducted the bioinformatics analysis and data visualization.

S.R.: contributed to the study’s conception, experimentation, data analysis, interpretation, helped in writing the manuscript.

S.R knocked out *SNCA* in WM983B cell line, performed lentiviral transduction, and validated RNA-seq by RT-PCR in WM983B and SK-MEL-28 cells.

SS: participated in the study design, knocked out the *SNCA* in SK-MEL-28 cells, and contributed to the interpretation of the results.

SNW: conception, project oversite, data analysis, wrote and edited the manuscript.

XY: Participated in the discussion about the genes that are differentially expressed in the KO cells.

## Figure legends

**Fig. S1.**
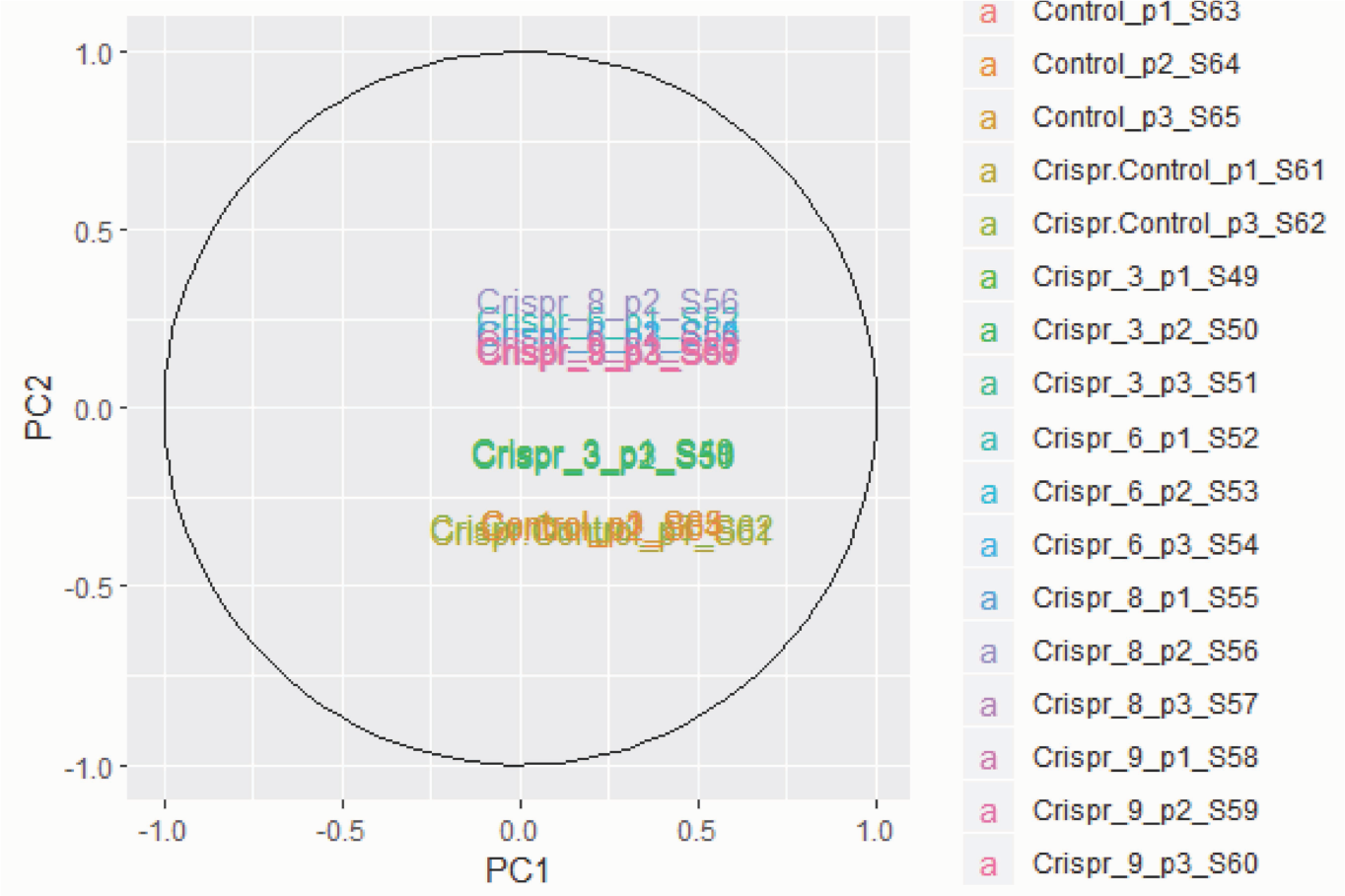
Principal component analysis (PCA) of RNA-seq samples. The principal components 1 (X-axis) and 2 (Y-axis) were used to coordinate the sample transcriptomes. The samples were colored based on their grouping information. The plot shows that the five control samples cluster together, the nine homozygous *SNCA*-KOs cluster together in a separate space, and three hemizygous *SNCA*-KOs form a third cluster in between the control and KO clusters.

**Fig. S2.**
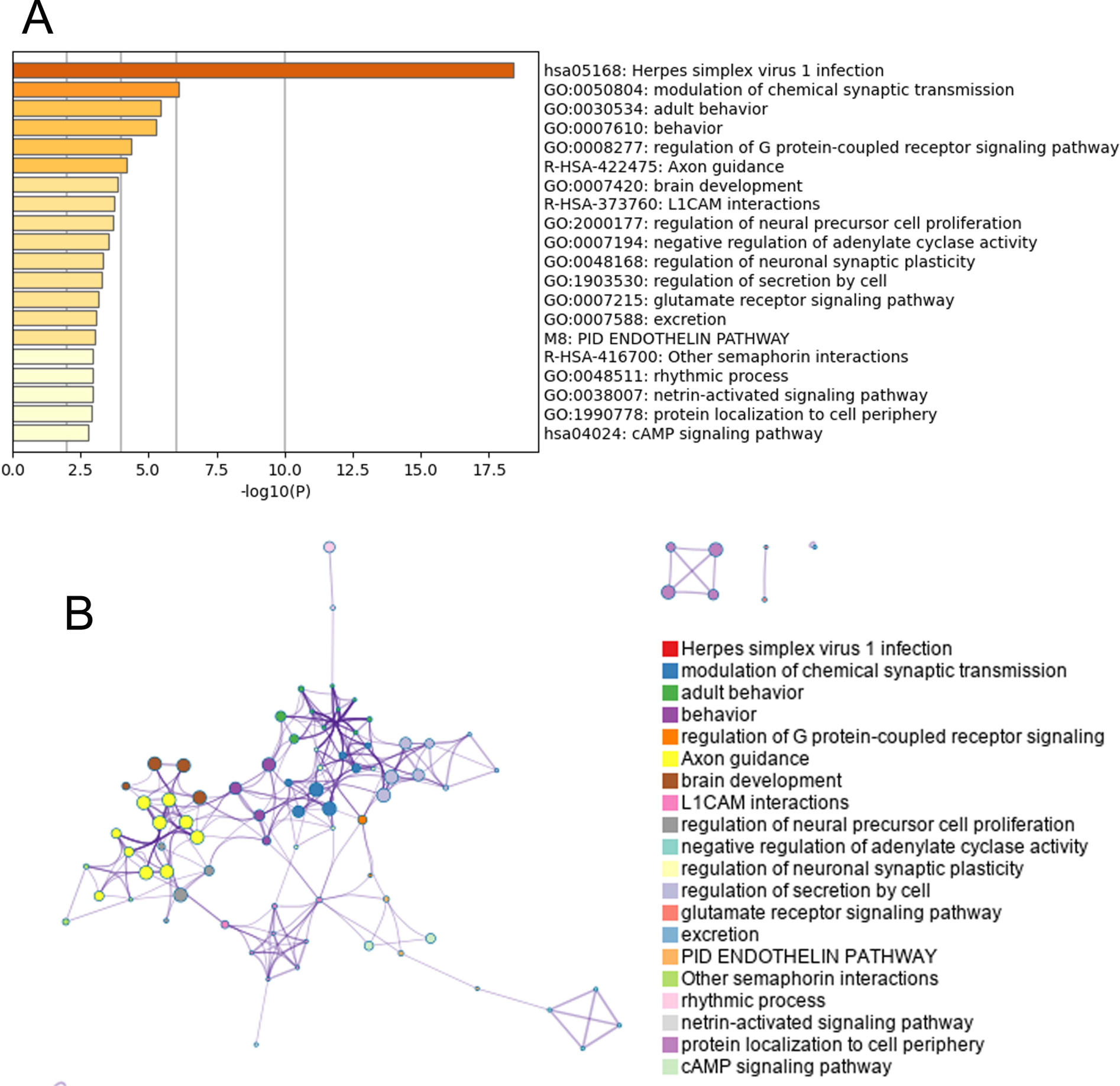
Gene ontology and network analyses. (**A**) Gene ontology enrichment analysis shows 20 enrichment clusters of down-regulated gene in *SNCA*-KO clones. (**B**) Enrichment network visualization of down-regulated genes in *SNCA*-KO clones. Both analyses conducted at http://metascape.org.

**Fig. S3.**
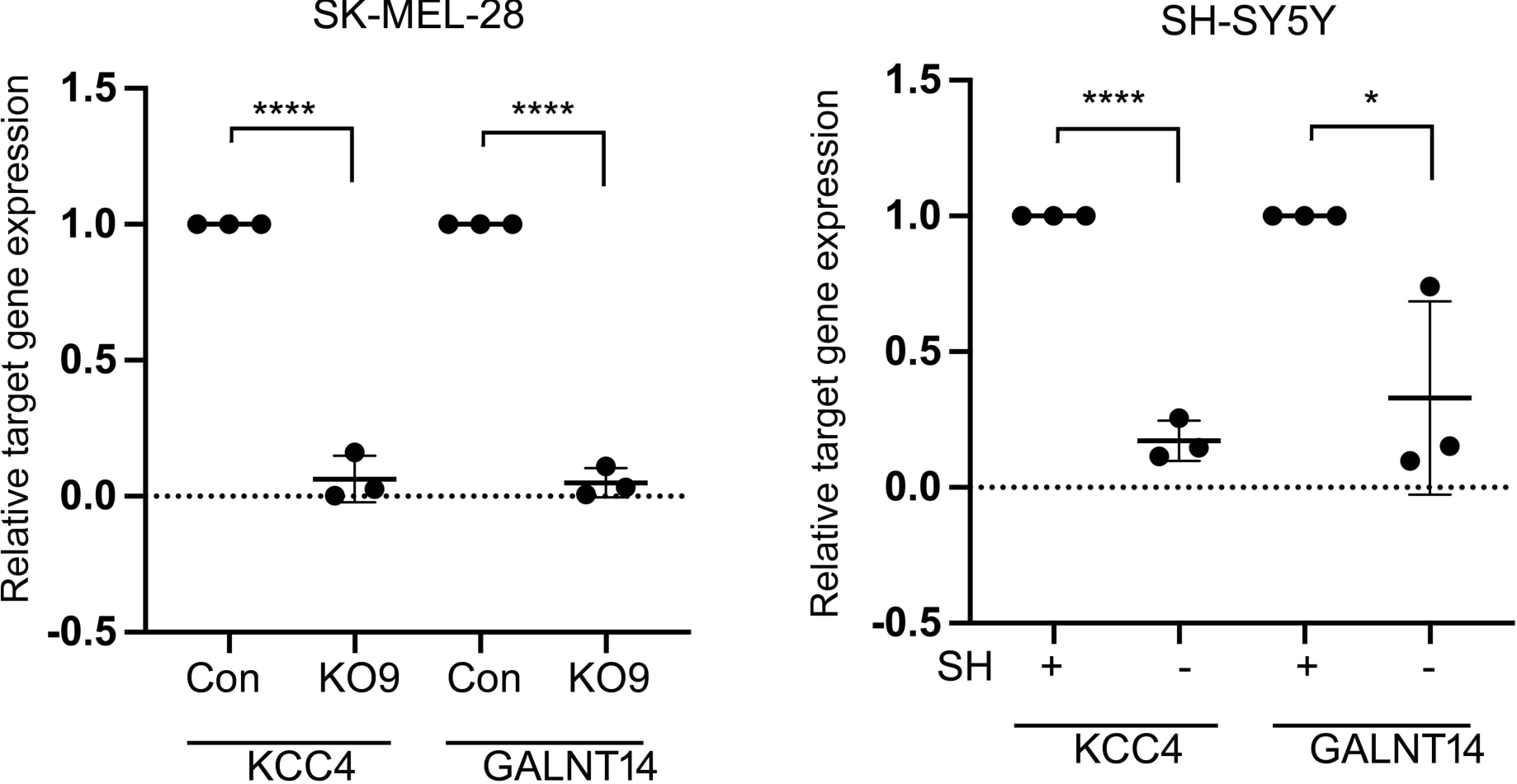
RT-qPCR validation of two down-regulated genes: KCC4 and GALNT14. These two transcripts had very low base mean values in control and KO cells (<100); nevertheless, we tested whether these transcripts were down-regulated by RT-qPCR. (**A**) Relative KCC4 and GALNT14 transcript levels in SK-MEL-28 control and KO clones. (**B**) Relative KCC4 and GALNT14 transcript levels in SH-SY5Y low (-αS) and high (+ αS) α-syn-expressing cells. Data are mean ± s.d. P-values were determined using a one-tailed Student’s t-test with equal variances (n=3). *, *p* < 0.05, ****, *p* < 0.0001.

**Fig. S4.**
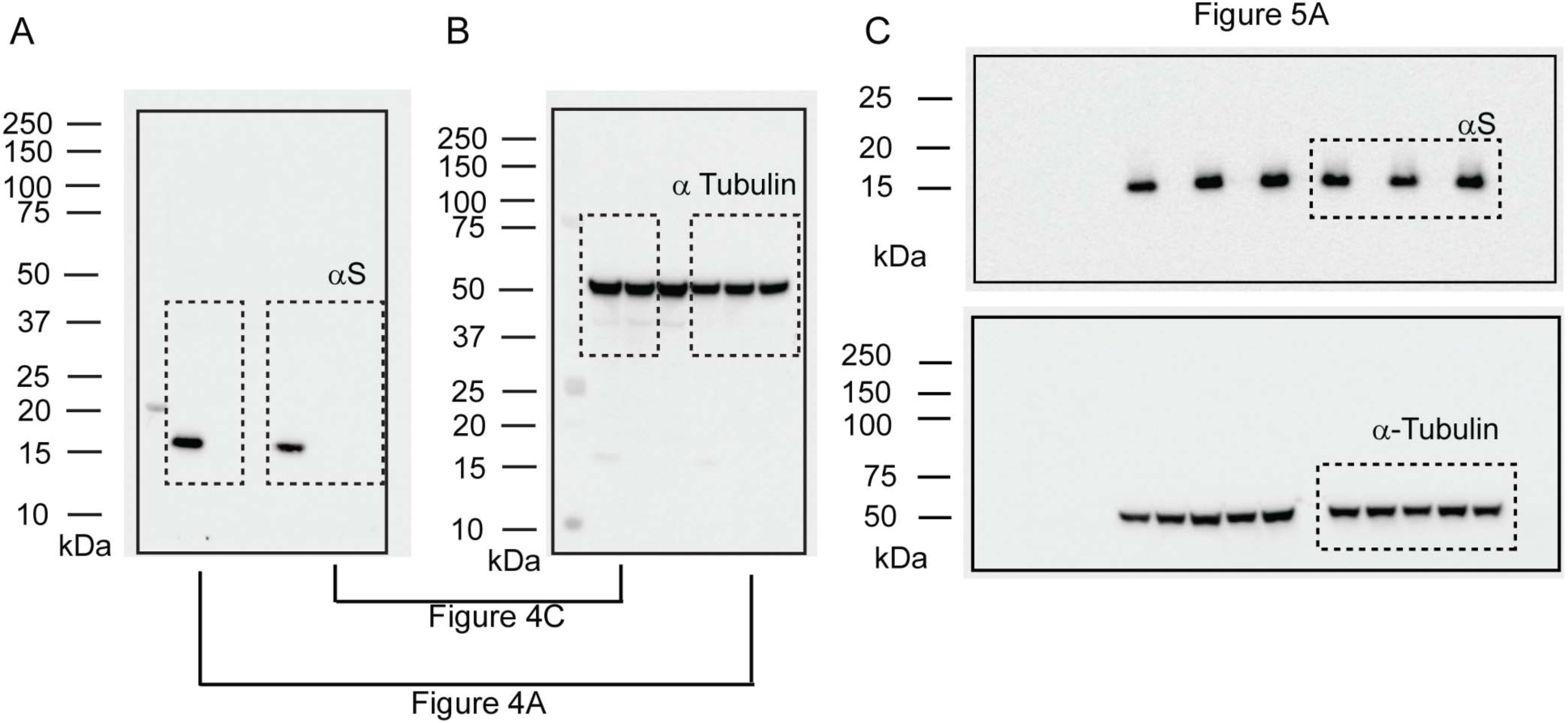
Uncropped blots. Uncropped blots to the indicated figures.

